# A Multimodal Deep Learning Framework for Predicting PPI-Modulator Interactions

**DOI:** 10.1101/2023.08.03.551827

**Authors:** Heqi Sun, Jianmin Wang, Hongyan Wu, Shenggeng Lin, Junwei Chen, Jinghua Wei, Shuai Lv, Yi Xiong, Dong-Qing Wei

## Abstract

Protein-protein interactions (PPIs) are essential for various biological processes and diseases. However, most existing computational methods for identifying PPI modulators require either target structure or reference modulators, which restricts their applicability to novel PPI targets. To address this challenge, we propose MultiPPIMI, a sequence-based deep learning framework that predicts the interaction between any given PPI target and modulator. MultiPPIMI integrates multimodal representations of PPI targets and modulators, and uses a bilinear attention network to capture inter-molecular interactions. Experimental results on our curated benchmark dataset show that MultiPPIMI achieves an average AUROC of 0.837 in three cold-start scenarios, and an AUROC of 0.994 in the random-split scenario. Furthermore, the case study show that MultiPPIMI can assist molecular simulations in screening inhibitors of Keap1/Nrf2 PPI interactions. We believe that the proposed method provides a promising way to screen PPI-targeted modulators.

## INTRODUCTION

Protein-protein interactions (PPIs) play a critical role in numerous cellular processes, such as cell proliferation, survival and apoptosis ^1–4^. Dysfunction in PPIs have been associated with various physio-pathologies including cancer development, infectious diseases and neurological disorders ^5^. Unlike traditional drug targets with active sites, which may share similarities with other regions of the proteome and cause unwanted off-target effects, the diversity of PPIs offers a better chance for developing selective and specific drugs ^6,7^. Therefore, modulating PPIs with small molecules is a promising strategy to treat diseases. However, identifying PPI modulators (inhibitors/stabilizers) is challenging due to the special structural properties of PPIs. For example, they have highly variable interfaces without well-defined binding pockets, and larger interface area (1000-2000 Å2) with high hydrophobicity ^6^. Based on these properties, PPI modulators are often large hydrophobic molecules with poor oral bioavailability and cell permeability. Despite this, many studies have shown that hotspot regions in PPIs, which contribute most to the binding affinity, can be targeted by compounds with lower molecular weight ^8–10^.

High-throughput screening is a conventional approach for identifying PPI modulators. Nevertheless, it has a low success rate compared to enzymes or individual protein receptors, due to the need for special considerations to identify interaction hotspots ^6^. In contrast, *in silico* screening of PPI modulators (PPIMs) can greatly narrow the search space for candidate modulators. Structure-based approaches, such as molecular docking and molecular dynamics simulations, are commonly used to screen large compound libraries against PPI partner proteins ^11^. However, finding novel scaffolds by docking is still challenging, mainly due to the large and featureless interfaces of PPI complexes ^12^.

Machine learning (ML) or deep learning (DL) has become a powerful tool for drug discovery, owing to the availability of large public databases ^13,14^. Among them, several databases focus on PPIMs, such as 2P2I-DB ^15^ which contains over 274 inhibitors that target 27 distinct PPIs, and iPPI-DB ^16^ which covers over 2378 modulators that interact with 46 PPI families. These datasets enable the application of data-driven methods for screening potential PPIMs. For example, Hamon *et al.* use DRAGON descriptors and Support Vector Machine (SVM) algorithm to identify novel PPI inhibitors based on 40 known ones ^17^. PPIMpred applies SVM with 11 molecular features to predict inhibitors for three oncogenic PPIs^18^. SMMPPI adopts a two-stage classification method to predict PPIMs based on molecular structure fingerprints ^19^. pdCSM-PPI utilizes a graph-based compound representation to facilitate the discovery of PPIMs for 21 different PPI targets ^20^. However, existing methods predict modulators based on similarity to known ones, while ignoring PPI target information and the interaction principles between PPI targets and modulators. Consequently. these approaches are not suitable for novel or under-studied PPI targets that lack sufficient known modulators. Therefore, it is necessary to develop generalizable approaches that can directly predict the interactions between any PPI target-modulator pairs.

Effective representations of compounds and proteins are essential for predicting PPI target-modulator interactions (PPIMI). DL-based approaches can effectively learn the representations (embeddings) from raw sequential or structural data of compounds and proteins, using expressive neural network architectures such as Transformer. For instance, Transformer-based protein pre-trained models such as ESM-2, TAPE and ProtTrans have demonstrated outstanding performance in various downstream tasks, including structure and function prediction for proteins ^21–23^. For compounds, pre-trained graph neural network (GNN) models can learn high-level representations of compounds from molecular graphs using self-supervised learning (SSL) ^24–27^. Graph SSL can be either contrastive or generative, depending on the supervision signals. Contrastive graph SSL learns the representation by contrasting with other graphs ^28–30^, while generative graph SSL focuses on reconstructing the original graph ^28,31^. On the other hand, physicochemical properties are also important for characterizing molecules, with examples including molecular weight, solubility and lipophilicity for compounds, and aliphaticity, hydrophilicity, and polarity for proteins. However, structural representations alone may not be sufficient to capture the complex interactions between PPI targets and modulators. Integrating multimodal representations for proteins and compounds can generate more comprehensive representations and enhance prediction accuracy. This has been demonstrated in DL-based models for compound-protein interaction (CPI) prediction ^32–34^. As such, the exploration of multimodal data holds promise in predicting PPIMI.

In this study, we propose a novel deep learning framework named MultiPPIMI to predict the interactions between PPI target and modulator (**Figure 1**). The model integrates multimodal representations of modulators and PPI targets, including structural embeddings and physicochemical properties, to comprehensively capture their characteristics. The model also uses bilinear attention networks to explicitly learn the inter-molecule interactions between PPI targets and modulators. The main contributions of this study are as follows:

- We construct a benchmark dataset that contains information on PPI targets, modulators, and their interaction labels.
- We are the first to predict PPI target-modulator interactions by formulating it as a binary classification problem, with the focus on screening PPI modulators for any given target.
- We experimentally validate the efficacy of multimodal representations and the bilinear attention mechanism in enhancing prediction accuracy and generalization capability across warm-start and cold-start scenarios.
- We show that combining MultiPPIMI with molecular simulations improves the hit rate for screening inhibitors of Keap1/Nrf2 PPI interactions.

**Figure 1.**
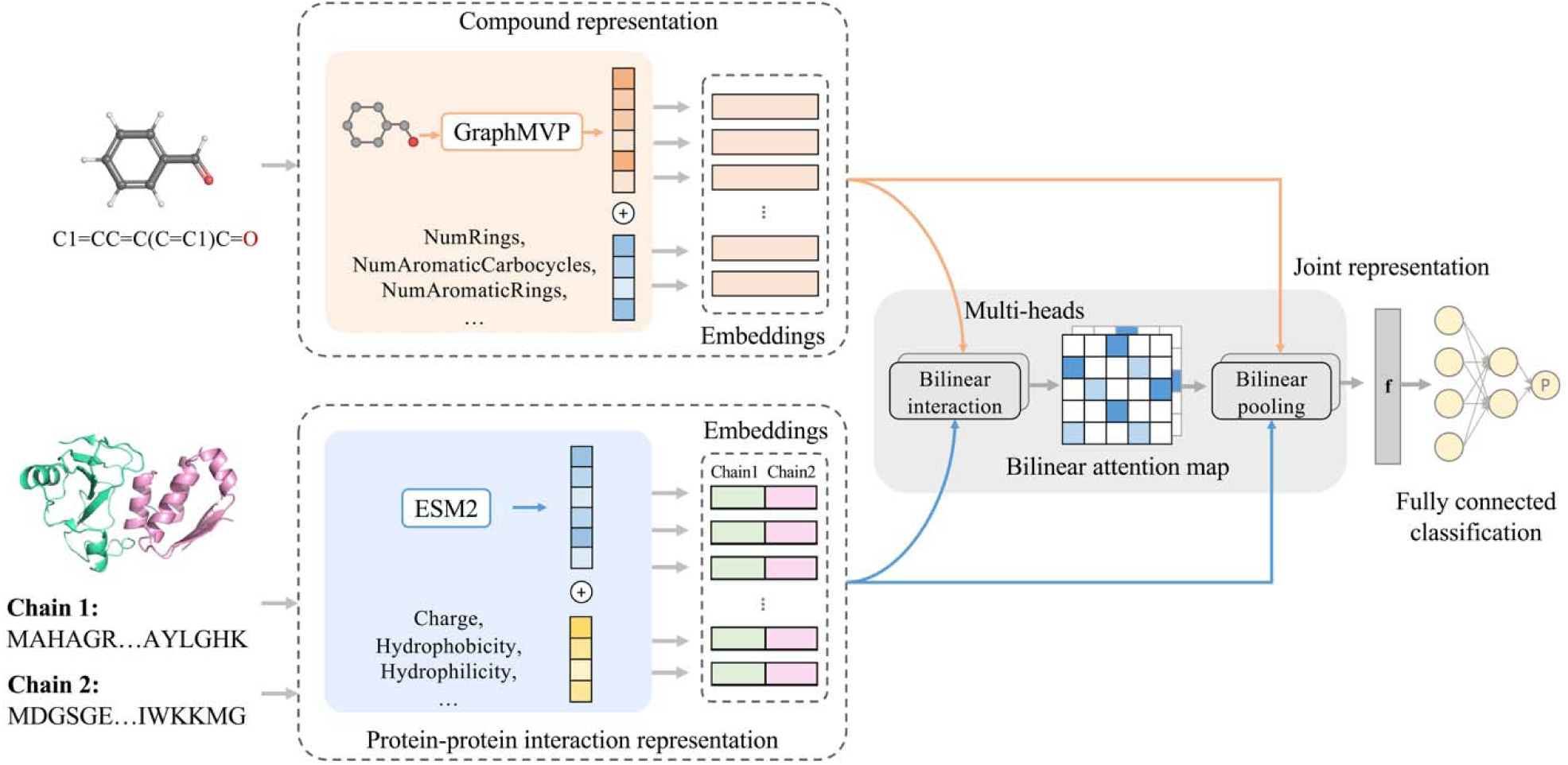
Overview of the MultiPPIMI framework. The modulator representation consists of the concatenation of pre-trained structural embeddings and physicochemical properties of compounds. The PPI representation is constructed by concatenating ESM2 embeddings and physicochemical features of two protein chains. The modulator and PPI representations are input into a bilinear attention network, enabling the learning of inter-molecular interactions. The joint representation **f** is decoded using a fully connected network to predict the probability *p* of PPI target-modulator interactions.

## METHODS

### Problem formulation

In PPIMI prediction, the objective is to predict the interaction between modulators and a PPI target. If the PPI target involves more than two partners, we randomly select two of them to represent the complex. The amino acid sequence of each partner is denoted as, where represents one kind of amino acid. Therefore, each PPI target is represented as. Regarding the modulator, we use the simplified molecular-input line-entry system (SMILES) representation to calculate physicochemical features, and convert it into a 2D molecular graph for structural embedding calculation. A molecular graph, defined as, consists of a set of vertices (atoms) denoted as V, and a set of edges (chemical bonds) denoted as E. The goal of PPIMI prediction is to train a model M that maps the joint feature representation space to an interaction probability score *p* ∊ [0, 1].

### Datasets

To evaluate our proposed model for PPIMI prediction, we construct a benchmark dataset based on a public database DLiP ^35^, which contains active and inactive small modulators of PPI targets curated from public databases and published literatures. The protein sequences of PPI partners are retrieved from the UniProt database ^36^. With strict data filtering procedures (Supporting Information, section 1), we retain 9,817 distinct modulators and 120 PPI targets. Notably, over 80% of the modulators have a pairwise similarity coefficient less than 0.6 (Figure S1) calculated via Tanimoto using ECFP4 fingerprints, indicating a low redundancy in our filtered dataset.

To construct a balanced PPIMI dataset, positive samples are defined as pairs of modulators and PPI targets that interact with each other. To generate negative samples, we adopt a similar approach as a previous study ^19^. For each PPI family, we assume all modulators that interact with the targets from the other PPI families are inactive. To avoid mis-assigning inactive modulators, we exclude all putative negative modulators that overlap with true active modulators. Negative samples are defined as pairs of PPI targets and putative inactive modulators. To address class imbalance, we perform under-sampling on the negative samples, resulting in a balanced dataset of 11,630 positive pairs and 11,630 negative pairs. Notably, we exclude experimentally validated inactive PPIMs from the original DLiP dataset to reduce noise, because many PPI targets do not have validated inactive PPIMs.

### The framework of MultiPPIMI

The MultiPPIMI framework is designed to directly model PPI-modulator interactions (**Figure 1**). It takes PPI-modulators pair as input and generates the probability score of their interaction. This framework consists of four main modules: modulator feature extraction, PPI feature extraction, interaction modelling, and classification. In the modulator feature extraction module, the structural embeddings and physicochemical properties are combined to represent each compound. Similarly, structural embeddings and physicochemical property features of two protein chains are concatenated to generate a PPI representation vector. To model the local interactions between modulators and PPI complexes, we employ a bilinear attention network which captures the intermolecular interactions and fuses the multimodal representations. Finally, a multilayer perceptron (MLP) is utilized as the classifier to predict the probability of interaction between a candidate modulator and a PPI target. **Compound representation.** Compound embeddings are computed using the Graph Multi-View Pre-training (GraphMVP) framework ^25^. This framework employs a GNN-based multi-view pre-training network to establish the connection between 2D topological views and 3D geometric views of small compounds. The input of GraphMVP is 2D molecular graph, where atoms and bonds are represented as nodes and edges, respectively. Even though the encoder takes only 2D molecular graphs as input, the resulting embeddings effectively incorporate the 3D modality. The output is a 300-dimensional embedding vector for each compound.

Additionally, physicochemical properties are considered as a complementary modality to the structural view. These features are computed using RDKit ^37^ (Table S1). A 1×10 feature vector is generated for each input compound.

### Protein-protein interaction representation

For PPI target representation, we combine the structural and physicochemical modality. The structural embeddings are obtained using the pre-trained protein language model ESM2, due to its success in various tasks ^38^. ESM2 is an unsupervised Transformer model trained on protein sequences from the UniRef database ^39^. Despite its simple masked language modeling objective during training, ESM2 effectively produces structural features by leveraging the diversity of protein sequences. In this study, we use the 150M ESM2 model as an encoder, generating 640-dimensional embeddings for each protein sequence.

Physicochemical properties are also important as they can influence protein types, structures, and functions. Since protein sequences consist of 20 unique amino acids, each with distinct chemical and physical properties, we utilize 19 physicochemical features for each amino acid to capture this diversity (Supplementary Table S2). Using the Pfeature tool ^40^, a 1×19 vector is generated for each protein sequence, representing the combined physicochemical properties of all its amino acids.

### Bilinear interaction networks

Bilinear attention networks are initially designed for visual question answering, a task involving multiple modalities ^41^. These networks can capture interactions between pairs of multimodal input channels, such as image and text, allowing for the integration of diverse modalities and facilitating the learning of effective interaction representations. Bilinear attention networks excel at capturing rich joint information in multimodal data while maintaining computational efficiency comparable to unitary attention mechanisms.

In this study, we use an adapted version of a bilinear attention network from a recent study on CPI prediction ^42^. This module serves to model non-covalent interactions between modulators and PPI targets. The bilinear attention module consists of two layers. The first layer is bilinear interaction map that captures pair-wise attention weights. It takes the representations of PPI targets and modulators as input, referred to as **H_P_ = [h_1p_, h_2p_, h_3p_, … h_Mp_]** and **H_c_ = [h_1c_, h_2c_, h_3c_, … h_Nc_]**, where M and N denote the dimensions of PPI and modulator representations, respectively. Then, a bilinear interaction map is constructed from these representations, resulting in a pairwise interaction matrix **I** ∊ ℛ^M×N^ for a single head:

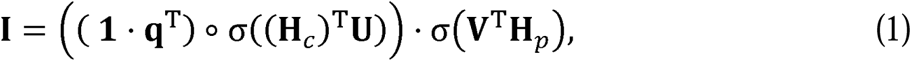

which defines the relationship between the learnable weight matrices **U** ∊ ℛ^D_c_×k^ and **V** ∊ ℛ^D_p_×k^, the learnable weight vector **q** ∊ ℛ^*k*^, the fixed all-ones vector **1** ∊ ℛ^*N*^, and the element-wise product ֯. Intuitively, the bilinear interaction maps the representations of PPIs and compounds to a common embedding space using the weight matrices **U** and **V**. The interaction is then learned by applying an element-wise product and the weight vector **q**.

The second layer is a bilinear pooling layer that extracts a joint representation **f**′ ∊ ℛ^*k*^ of the PPI-modulator pair from the interaction map **I**. Specifically, the *k*th column of **f**′ is calculated as follows:

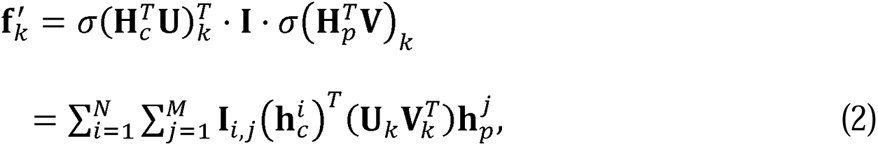

where the **U** and **V** are shared with the interaction map layer. Subsequently, a feature map is generated by applying sum pooling to the joint representation vector:

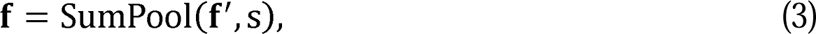

where the *sumPaal*(.) function refers to a non-overlapping 1D sum pooling operation with a stride *s*. In our approach, the multi-head version of the single pairwise interaction is employed by summing the representation vectors from individual heads. Each head is associated with a specific weight vector **q**, while the weight matrices **U** and **V** are shared across all heads.

### Classification module

The feature map **f** obtained from the bilinear attention module is passed through fully connected layers with the ReLU activation function, as depicted below:

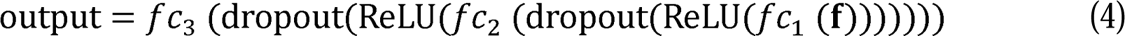

Subsequently, a softmax layer is added after the fully connected layers for the final prediction:

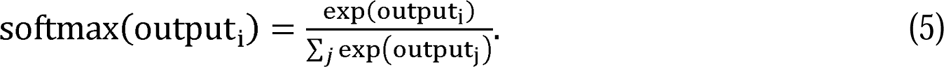

The output value represents the probability of interaction between a PPI-modulator pair.

### Different pre-training strategies of compound structural encoder

This study compares three self-supervised learning tasks for compound encoder GraphMVP. The original pre-training objective is defined as follows:

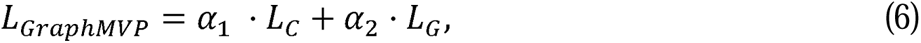

where *α*_1_, *α*_2_ are weighting coefficients. *L_C_* represents the contrastive loss, which treats 2D-3D pairs of the same molecule as positive and negative otherwise. *L_G_* is the reconstruction loss used for generating 3D conformers from the corresponding 2D topology.

The original pre-training strategy focuses on enhancing the 2D representation by leveraging 3D information (Supporting Information, section 2). To further exploit the 2D topology, we additionally include 2D SSL tasks in the original objective as illustrated in ^25^. We compare two main categories of 2D SSL tasks: generative and contrastive. This results in two variants of *L_GraphMVP_,* namely GraphMVP-G and GraphMVP-C, with the following objectives:

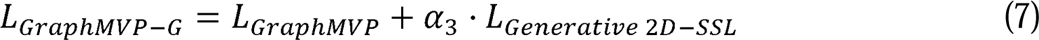

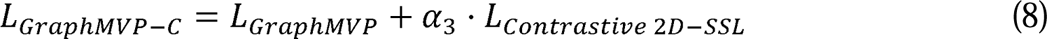

### Evaluation strategies and metrics

We use four data split strategies to evaluate the performance of PPIMI prediction models on the benchmark dataset. These strategies reflect different levels of generalization:

- S1: Random split setting, where 5-fold CV is performed on PPI-modulator pairs.
- S2: Modulator cold-start setting, where 5-fold CV is conducted on modulators. The unique compounds are clustered into five groups based on their chemical structure similarity using the Butina clustering algorithm with ECFP4 fingerprints and a clustering radius of 0.8.
- S3: PPI cold-start setting, where 5-fold CV is conducted on PPI targets.
- S4: Cold pair setting, where 5-fold CV is conducted on both compounds and PPI targets. In this scenario, both the testing compounds and testing targets are unseen in the training set.

The performance of binary classification is evaluated using area under the receiver-operating characteristic curve (AUROC), area under the precision-recall curve (AUPR), sensitivity, precision, and specificity.

### Implementation

MultiPPIMI is implemented using Python 3.7.15 and PyTorch 1.8.0 ^43^. The model is trained using cross-entropy loss with a batch size of 64, and the Adam optimizer with a learning rate of 0.0005. The model is trained for a maximum of 500 epochs. The best-performing model is selected based on the epoch that yielded the highest AUROC score on the validation set. All hyperparameters related to the GraphMVP compound encoder and the bilinear attention module are set to their default values as specified in the original papers. The hyperparameters of batch size, learning rate, and epochs are fine-tuned under the random-split setting and remain consistent across all evaluation scenarios.

### Baselines

We compare the performance of MultiPPIMI with four baseline models. Three baselines are shallow machine learning classifiers: Support Vector Machine (SVM) ^44^, Random Forest (RF) ^45^, and XGBoost ^46^. The fourth one is a neural network classifier: Multilayer Perceptron (MLP). All classifiers use the same input features: the concatenation of ECFP4 fingerprint ^47^ for compounds and PseAAC ^48^ for PPI partners. We finetune these classifiers using the AUROC on the validation set under the random split setting. The hyperparameter search grid and the optimal values for each classifier are listed in Table S3.

## RESULTS AND DISCUSSION

### Performance evaluation

We compare MultiPPIMI with four baseline models using the four different settings (S1-S4): Random Forest (RF), XGBoost, Support Vector Machine (SVM), and Multilayer Perceptron (MLP). S1 is a random split setting with minimal domain shift, where both the testing compounds and PPI targets may appear in training set. S2-S4 are cold-start settings that are more realistic and challenging, which introduce unseen modulators, PPI targets, or both in the test set, respectively. Table 1 shows the performance comparison.

**Table 1.**
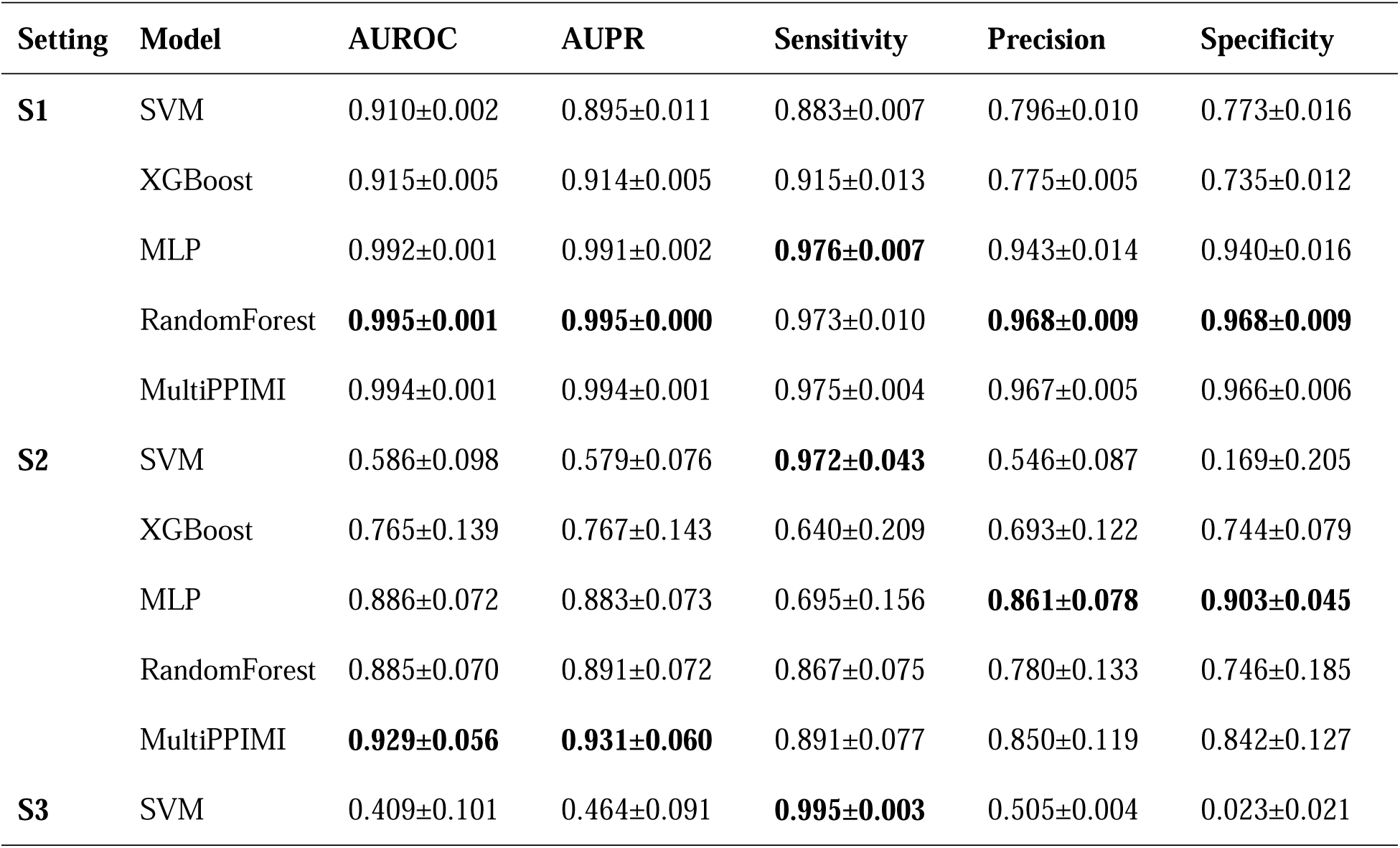
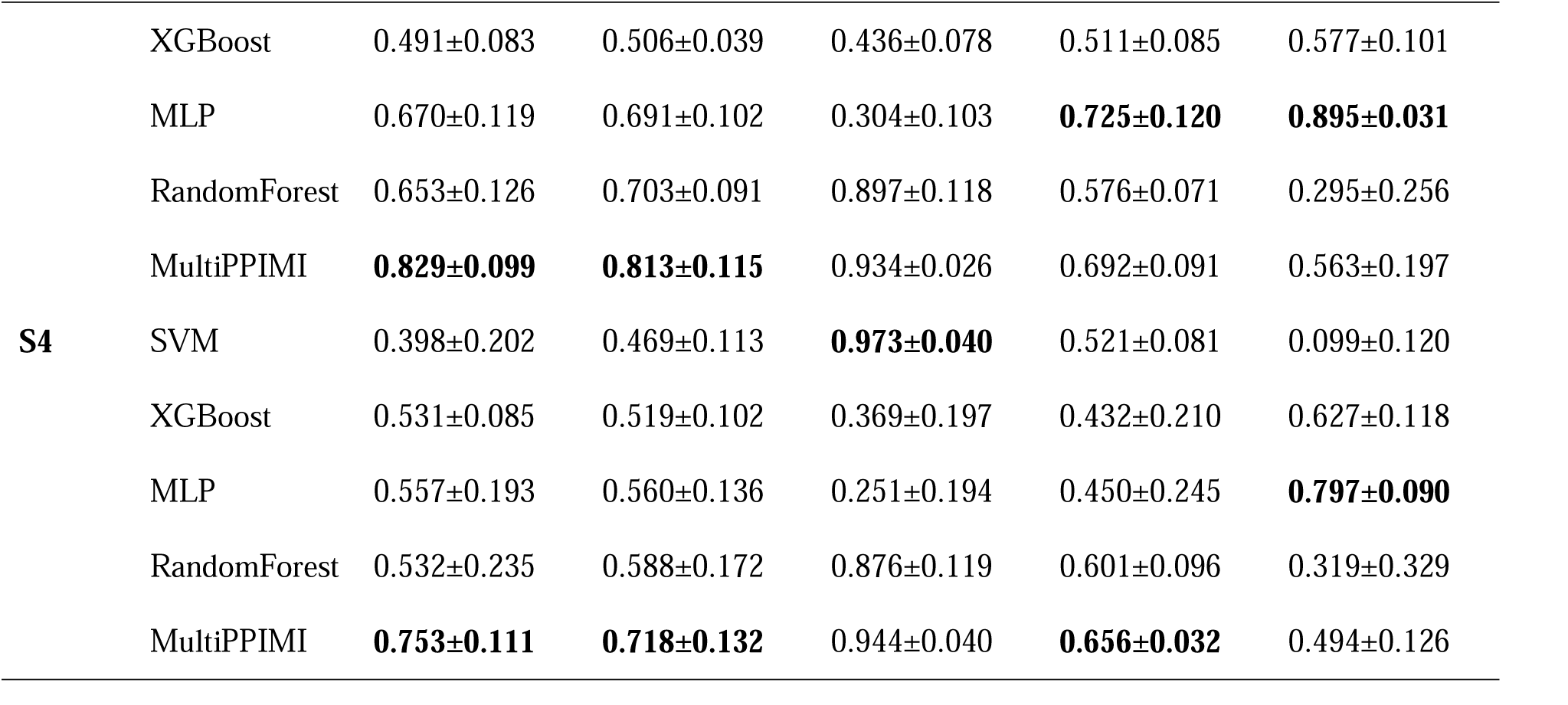
Performance comparison of different data split settings over 5-fold cross validation (mean±standard deviation). The best results for each task are marked in bold.

Under setting S1, MultiPPIMI, RF, and MLP achieve comparable and high performance (AUROC > 0.99). However, this may reflect bias and overfitting rather than real-world performance, as a previous study on CPI prediction ^49^ reports that models may rely on compound patterns to make predictions due to potential ligand bias in the dataset, rather than learning interaction rules. PPIMI prediction is analogous to CPI prediction, so we conduct further evaluations in three cold-start settings (S2-S4) to mitigate the overoptimism in random-split performance. These scenarios ensure that prediction on test data cannot rely only on the features of known modulators or PPI targets. All models show a significant performance drop compared to setting S1, but MultiPPIMI consistently outperforms the baselines in terms of AUROC and AUPR. Notably, MultiPPIMI exhibits higher average AUROC and AUPR in setting S2 than in setting S3, indicating that testing on new modulators is generally easier than on new PPI targets. This observation can be attributed to the larger number of modulators available in the training dataset than targets. In the most challenging scenario S4, the AUROC gap between MultiPPIMI and the baselines is the largest, demonstrating that MultiPPIMI can better learn the general principles of PPIMI and is more robust in real-world-alike conditions.

### Ablation study

In this study, we conduct an ablation study to assess the contribution of each main component in MultiPPIMI. We investigate the influence of each feature modality by individually removing them from the input (**Figure 2a**). The results reveal that the GraphMVP and ESM2 embeddings play a crucial role across all four tasks, showing the largest contribution in terms of AUROC. Moreover, incorporating physicochemical properties positively impacts cold-start predictions (S2-S4) but exhibits minimal effect on the warm-start setting (S1). This suggests that for simpler tasks involving in-distribution generalization, incorporating overly complex features may be unnecessary.

**Figure 2.**
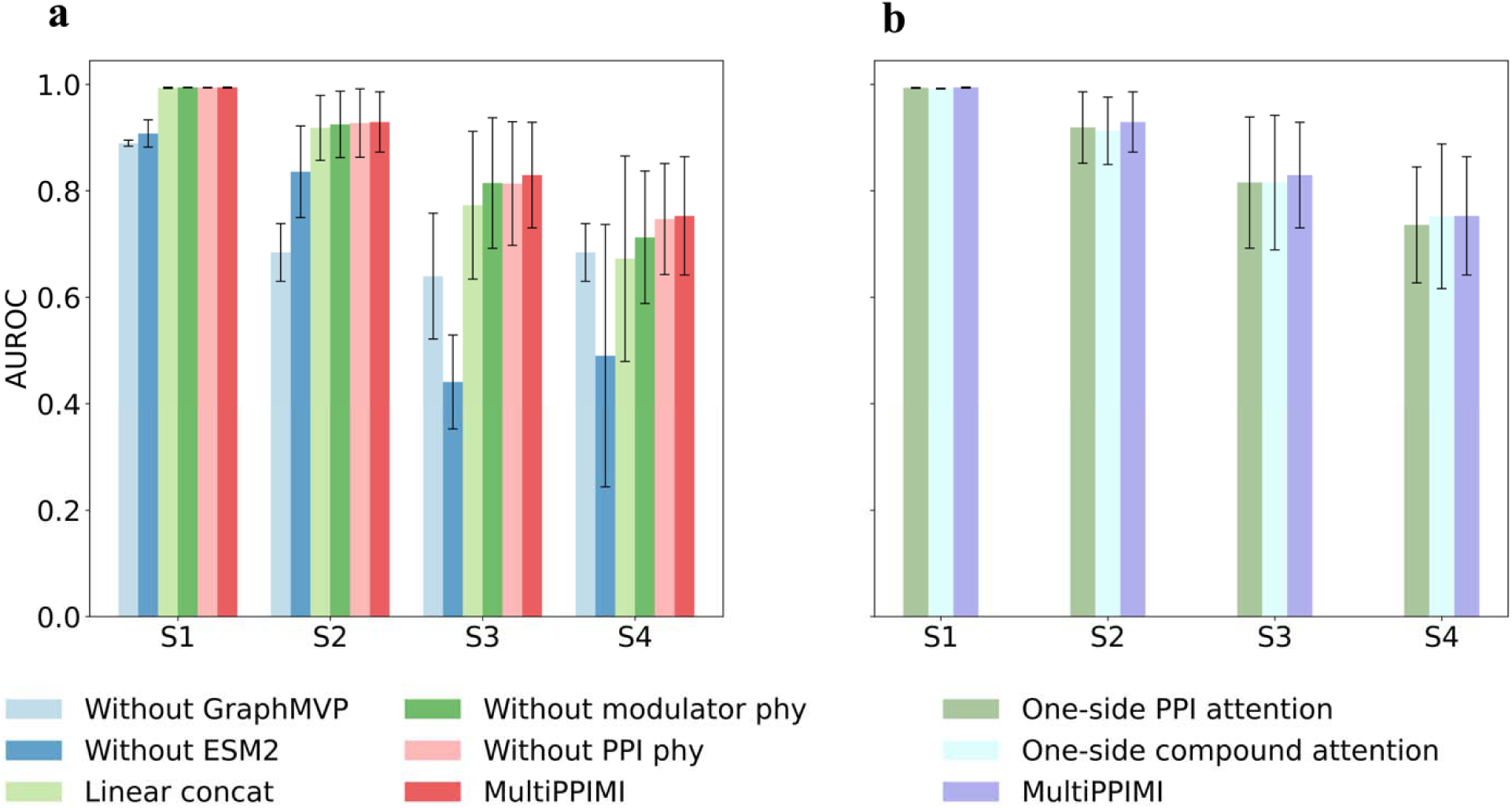
Ablation study with 5-Fold cross-validation across different data split settings. (a) Ablation study on the key components of MultiPPIMI. ‘phy’ denotes physicochemical features. (b) Ablation study on bilinear attentio module.

Additionally, we investigate the effectiveness of bilinear attention by comparing three variants of MultiPPIMI, each differing in the calculation of joint representations between modulators and PPI targets: one-side modulator attention, one-side PPI attention, and linear concatenation (**Figure 2a** and **Figure 2b**). The one-side attention approach, inspired by a study on prediction of CPI ^50^, attends solely to the sub-patterns within modulator or PPI representations. We replace the bilinear attention in MultiPPIMI with one-side attention to create the two variants. Another variant uses linear concatenation, which simply combines modulator and PPI representations. The results demonstrate that bilinear attention is the most effective method for capturing interaction information in PPIMI prediction. Interestingly, the differences among the four variants are negligible under S1 but more significant under S2-S4, suggesting that bilinear attention is particularly valuable for enhancing cold-start generalization.

### Comparison of pre-training strategies to learn compound embeddings

We investigate the impact of different self-supervised learning (SSL) tasks on compound embeddings by employing various SSL tasks to pre-train GraphMVP on the GEOM dataset ^51^. **Table 2** shows the AUROC scores of MultiPPIMI and its variants that use different SSL tasks for the GraphMVP encoder. The variants are: no pre-training; GraphMVP, the original pre-training task that learns the correspondence between 3D geometry and 2D topology; GraphMVP-C, which adds a 2D contrastive SSL task to GraphMVP; and GraphMVP-G, which adds a 2D generative SSL task to GraphMVP. The 2D SSL acts as a regularizer to extract more 2D topological information in additional to 3D view.

**Table 2.** Comparison of AUROC for MultiPPIMI and its variants using different pre-training tasks for compound embeddings. The best result for each task is marked in bold.

|  | **S1** | **S2** | **S3** | **S4** | **Avg** |
| --- | --- | --- | --- | --- | --- |
| **GraphMVP-C** | **0.994±0.001** | **0.929±0.056** | **0.829±0.099** | 0.753±0.111 | **0.876** |
| **GraphMVP-G** | 0.992±0.001 | 0.916±0.068 | 0.828±0.104 | **0.764±0.119** | 0.875 |
| **GraphMVP** | 0.994±0.001 | 0.920±0.061 | 0.811±0.129 | 0.730±0.182 | 0.864 |
| **Without pre-train** | 0.991±0.001 | 0.913±0.058 | 0.822±0.113 | 0.751±0.140 | 0.869 |

On average, GraphMVP-C and GraphMVP-G outperform GraphMVP and no pre-training. This observation suggests that different modalities complement each other during the pre-training stage, which is consistent with previous findings that multimodal pre-training can enhance model generalization compared to single-modal pre-training ^33^. However, GraphMVP does not perform better than no pre-training, which suggests that the contribution of pre-training to PPIMI prediction depends on the choice of SSL tasks. Since GraphMVP-C outperforms the other variants on average, we choose it as the pre-training strategy in our final model.

### Parameter sensitivity analysis

We analyze the impact of three key parameters on the performance of MultiPPIMI: the number of attention heads, the hidden dimension, and the kernel size in the bilinear attention module. The number of attention heads varies from 1 to 10, the hidden dimensions range from 128 to 1024, and the kernel size varies from 2 to 10. **Figure 3** illustrates that MultiPPIMI are robust to these parameters, with only minor fluctuations observed. Therefore, we retain the default parameters as stated in the original paper ^42^: 2 attention heads, 512 hidden dimensions, and a kernel size of 3. Moreover, we find that multi-head attention consistently outperforms single head attention. This can be attributed to the fact that multi-head attention can model the interaction between modulators and PPI targets from multiple feature spaces, which is consistent with the diverse non-covalent interaction types observed between atoms and amino acids, such as hydrogen bonds and salt bridges.

**Figure 3.**
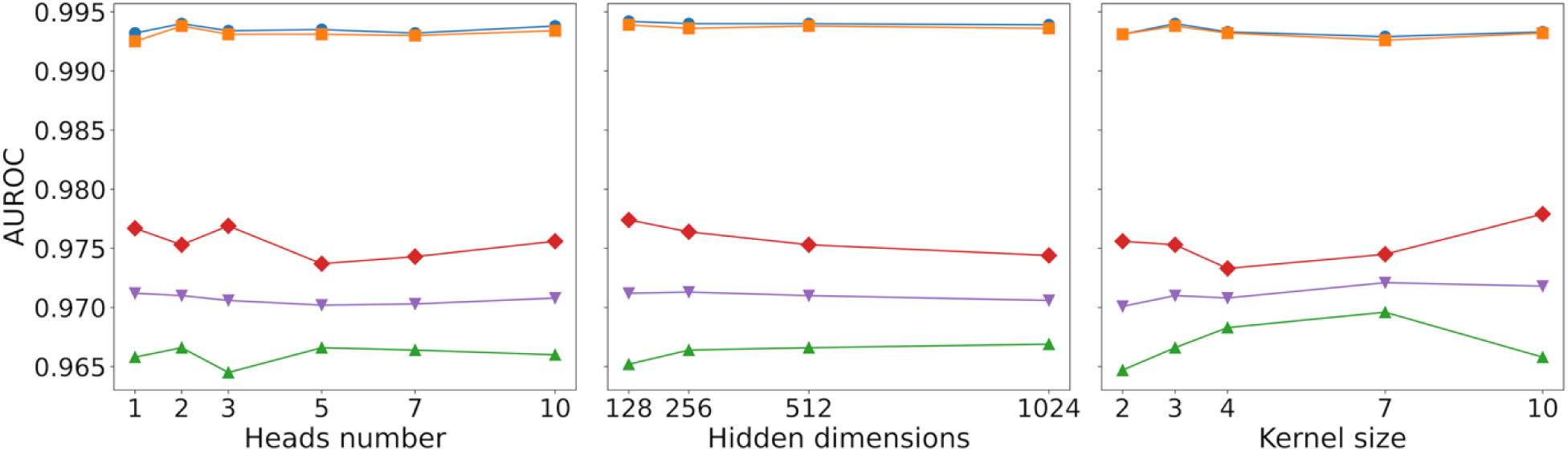
Parameter sensitivity analysis. Key parameters of MultiPPIMI are tested with 5-fold cross-validation under random split setting.

### MultiPPIMI-based screening for protein-protein interaction inhibitors

Due to the low hit rate of the primary screening of small molecular libraries against PPI targets, MultiPPIMI predictions can be used as a starting point for further virtual screening PPIs inhibitors ^52,53^. The Kelch-like ECH-associated protein 1/nuclear factor erythroid 2-related factor 2 (Keap1/Nrf2) PPI is an important target for the regulation of antioxidant defenses and has been relevant to a wide range of diseases including neurodegenerative, autoimmune and metabolic disorders ^53,54^.

We used the newly synthesized DLiP-PPI library ^35^ as the virtual screening library, which contains 15,074 compounds after deduplication and is independent of the training dataset of MultiPPIMI. We standardized all the candidate compounds by RDKit, and predicted their probability of interaction with Keap1/Nrf2 by MultiPPIMI. The hit compounds (probability > 0.7) were converted to 3D structures by RDKit and added AMBER ff14SB force field, AM1-BCC charges and hydrogen atoms by Chimera, before saving as mol2 files. Then we evaluate the drug-like space of the active inhibitors of Keap1/Nrf2 and the MultiPPIMI-predicted inhibitors, by mapping the uniform manifold approximation and projection (UMAP) plots ^55^ of MACCS fingerprints ^56^ and Ultrafast Shape Recognition with CREDO Atom Types (USRCAT) fingerprint^57^. As shown in UMAP plots (**Figure 4a** and **Figure 4b**), the hit molecules of MultiPPIMI share chemical space with the active inhibitors of Keap1/Nrf2 PPI target. We further performed molecular docking-based virtual screening on the hit compounds of MultiPPIMI and the reference compound using the UCSF DOCK6.9 program (Supporting Information, section 3) ^58,59^. The crystal structure of Keap1 protein (PDB ID: 4XMB) was selected for molecular docking ^60^. We selected the compounds with better docking scores than the reference compound, and visually inspected them by PyMOL ^61^. As shown in **Figure 4c**, the hit and reference compounds bind to the binding pocket formed by residues Tyr334, Arg380, Thr338, Arg415, Ile461, Arg483, Gln555, Ser555, and Tyr572. The hit compounds exhibit stronger binding affinity than the reference compound. Moreover, we overlapped the binding poses of the reference and hit compounds (**Figure S2)**. There is an overlap between the hit compound and the reference compound, and the hit compound better fits and merges into the interior of the pocket. Our results suggest that deep learning tools can complement molecular docking for screening PPI inhibitors by capturing the implicit relationships of PPI-modulator interactions, which provides a different perspective from the physical rule-based methods.

**Figure 4.**
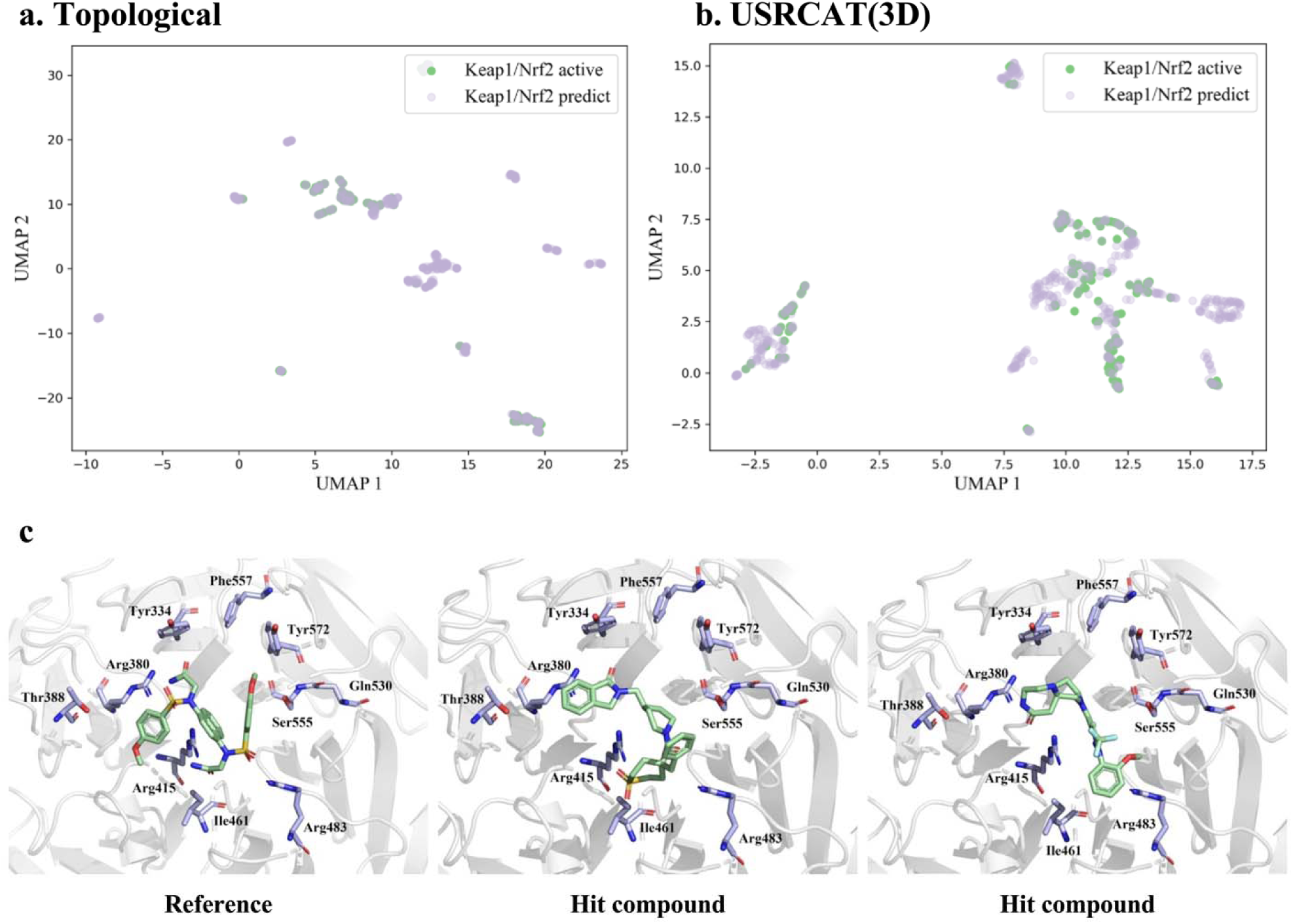
The UMAP visualization of the distribution of (a) MACCS fingerprint and the (b) USRCAT fingerprint in two-dimension space (violet: predicted inhibitors; green: active inhibitors). (c) Binding pose of the Keap1 protein with the reference compound (PDB ID: 4XMB) and binding poses of the hit compounds using docking.

## CONCLUSION

In this study, we construct a benchmark dataset of PPI-modulator interactions from public databases and propose MultiPPIMI, a deep learning framework for predicting the interactions between PPI target-modulator pairs. Our model integrates multimodal representations of modulators and PPI targets, including structural embeddings and physicochemical properties, to capture the diversity of both entities. Moreover, our model employs a bilinear attention network to learn the local interactions between PPI targets and modulators, which enables the extraction of the generalized interaction rules.

We conduct systematic evaluations to demonstrate the superiority of MultiPPIMI over the baseline model, especially in challenging cold-start scenarios that are frequently encountered in real-world applications. Furthermore, our results highlight the importance of multimodal pre-training of compound embeddings in enhancing PPIMI prediction, particularly in achieving robust out-of-distribution generalization. Case study demonstrate the potential of deep learning tools to facilitate molecular docking for virtual screening of PPI inhibitors. Unlike conventional physical rule-based methods, deep learning tools can capture the implicit relationships of PPI-modulator interactions in a data-driven manner, which provides a different perspective for identifying novel PPI inhibitors. However, MultiPPIMI concatenates features of both protein chains to represent PPI targets in high dimension, which has two limitations (i) it may increase computational cost as the training set grows, and (ii) it may overlook hotspot or interface information that are crucial for PPI modulation and model interpretability. In future research, we will explore more effective feature fusion strategies that can reduce dimensionality while preserving essential information. Additionally, we will exploit 3D structural information of PPI targets and integrating MultiPPIMI with deep molecular generative models ^62,63^ to discover novel PPI modulators.

## ASSOCIATED CONTENT

### Data Availability Statement

All the source code, software, and datasets in this publication is freely available under an MIT license at https://github.com/sun-heqi/MultiPPIMI. The benchmark dataset is stored in the directory “data”, the four data split scenarios are stored in “data/folds”.

### Supporting information

The following file is freely available.

Supporting Information: (1) Pre-processing of the DLiP curation dataset, (2) GraphMVP pre-training task, (3) Docking protocol; Figure S1 illustrates similarity of modulator pairs; Table S1 and Table S2 summarizes the physiochemical properties used by MultiPPIMI for small molecules and proteins, respectively; Table S3 lists the hyperparameter of the baseline classifiers (PDF)

### Author Contributions

Heqi Sun: conceptualization, formal analysis, writing original draft. Jianmin Wang: conceptualization, data analysis, writing review and editing. Hongyan Wu: supervision. Shenggeng Lin: writing review and editing. Junwei Chen: writing review and editing. Jinghua Wei: data analysis. Shuai Lv: data analysis. Yi Xiong: supervision, resources, writing review and editing. Dong-Qing Wei: supervision, resources, writing review and supervision.

### Notes

The authors declare no competing financial interest.

## ACKNOWLEDGMENTS

This work was supported by the National Natural Science Foundation of China [Nos. 32070662 and 61832019].

## For Table of Contents Only

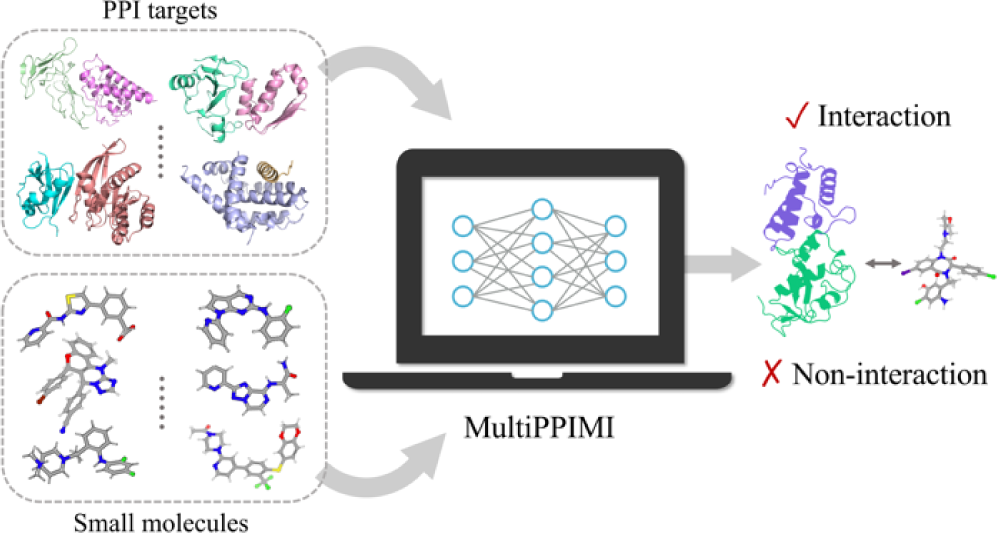

