## Supporting information for "A Multimodal Deep Learning Framework for Predicting PPI-Modulator Interactions"

**1 Pre-processing of the DLiP curation dataset**

The original DLiP curation dataset ^1^ undergoes strict filtering procedures before being used to construct our benchmark dataset. Firstly, the ‘Uncategorized’ and ‘AR/unknown partner’ PPI targets are excluded from the dataset. Additionally, certain PPI targets derived from the Timbal database ^2^ have broad categories, such as ‘Integrins’, ‘Bcl-2 and Bcl-XL with BAX; BAK and BID’, ‘Tubulin dimer’. Except for ‘Integrins’ which can be traced back to ChemBL ^3^, the original PPI targets for the remaining targets that cannot be tracked are therefore removed. To obtain more specific information, ChemBL database (V32) is used to trace the original PPI targets of ‘Integrins’. Activity values $\leq10\mu M$, as specified in the literature ^1^; data for the ‘Homo Sapiens’ species are retained. For the PPI targets corresponding to ‘BCL2-Like_BAX’, they are traced from the IPPI-DB database ^4^. PPI targets with composite-name like ‘BetaCatenin/Tcf4 and Tcf3’ are split into separate targets, resulting in duplicated DLiP IDs in the dataset. Finally, any duplicated pairs of PPI targets and small molecules are removed from the dataset.

**2 GraphMVP pre-training task**

The pre-training of GraphMVP ^5^ involves two types of self-supervised learning (SSL) tasks: contrastive SSL between 2D and 3D views and generative SSL between 2D and 3D views.

For contrastive SSL, pairs $\left( x,y \right)$ consisting of the 2D and 3D views from the same molecule are considered as positive, and negative otherwise. The objective function for contrastive SSL is based on an energy-based model with noise contrastive estimation (EBM-NCE):

$$\begin{aligned} \begin{aligned} L_{C}=&-\frac{1}{2}\mathbb{E}_{p\left( y \right)}\left[ \mathbb{E}_{p_{n}\left( x | y \right)}\log\left( 1-\sigma\left( f_{x}\left( x,y \right) \right) \right)+\mathbb{E}_{p\left( x | y \right)}\log\sigma\left( f_{x}\left( x,y \right) \right) \right]\#\# \\ &-\frac{1}{2}\mathbb{E}_{p\left( x \right)}\left[ \mathbb{E}_{p_{n}\left( y | x \right)}\log\left( 1-\sigma\left( f_{y}\left( y,x \right) \right) \right)+\mathbb{E}_{p\left( y,x \right)}\log\sigma\left( f_{y}\left( y,x \right) \right) \right], \\ & \end{aligned}\#\left( 1 \right) \end{aligned}$$

where $p_{n}$ denotes the noise distribution and $\sigma$ represents the sigmoid function.

The generative SSL aims to learn a robust 2D molecular representation capable of reconstructing its corresponding 3D conformer, and vice versa. The objective function for generative SSL is based on variational representation reconstruction:

$$\begin{aligned} \begin{aligned} L_{G}=&\frac{1}{2}\left[ \mathbb{E}_{q\left( z_{x} | x \right)}\left[ \left\| q_{x}\left( z_{x} \right)-SG\left( h_{y} \right) \right\|^{2} \right]+\mathbb{E}_{q\left( z_{y} | y \right)}\left[ \left\| q_{y}\left( z_{y} \right)-SG\left( h_{x} \right) \right\|_{2}^{2} \right] \right] \\ &+\frac{\beta}{2}\cdot[KL(q\left( z_{x} | x \right)\left\| p\left( z_{x} \right) \right)+KL\left( q\left( z_{y} | y \right)\left\| p\left( z_{y} \right) \right) \right],\# \end{aligned}\#\left( 2 \right) \end{aligned}$$

where SG is the stop-gradient operation.

The overall objective function for pre-training GraphMVP combines both contrastive SSL and generative SSL:

$$\begin{aligned} L_{GraphMVP}=\alpha_{1}\cdot L_{C}+\alpha_{2}\cdot L_{G},\#\left( 3 \right) \end{aligned}$$

where $\alpha_{1}$ and $\alpha_{2}$ are weighting coefficients.

**3 Docking protocol**

Data: The hit compounds were converted to 3D structures by RDKit. Each molecular structure was added AMBER ff14SB force field, AM1-BCC charges and hydrogen atoms by Chimera, before saving as a mol2 file.

The X-ray structure of the target was downloaded from the RCSB Protein Data Bank ^6,7^, and the 3D structure of Keap1 protein (PDB ID: 4XMB) was used for docking calculations ^8^. The protein structure was assigned the AMBER ff14SB force field and AM1-BCC charges ^9,10^, and the DMS tool in Chimera was utilized to generate the molecular surface of the receptor by using probe atoms with a radius of 1.4 Å ^11^. The binding sites of 41P (2,2'-(naphthalene-1,4-

diylbis(((4-methoxyphenyl)sulfonyl)azanediyl))diacetamide) in the X-ray structure were used in this docking study. For each site, the sphgen module was performed to generate spheres filling with active sites, and the grid module generated grid files for grid-based energy evaluation. The DOCK6.9 ^12,13^ program was used to perform semi-flexible docking. The van der Waals and electrostatic interactions were acquired between the ligand and the binding site, and the grid scores were calculated. Afterwards, clustering analysis (RMSD threshold set to 2.0 Å) was performed to obtain the best scored pose. For the other parameters, the default values were used.

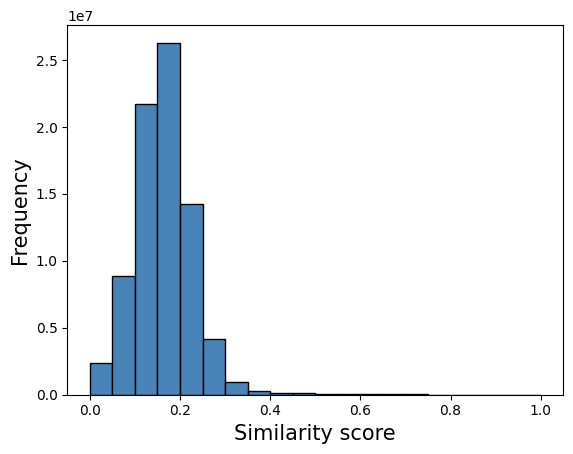

**Figure S1.** Tanimoto similarity distribution considering all the possible unique pairs of small molecules.

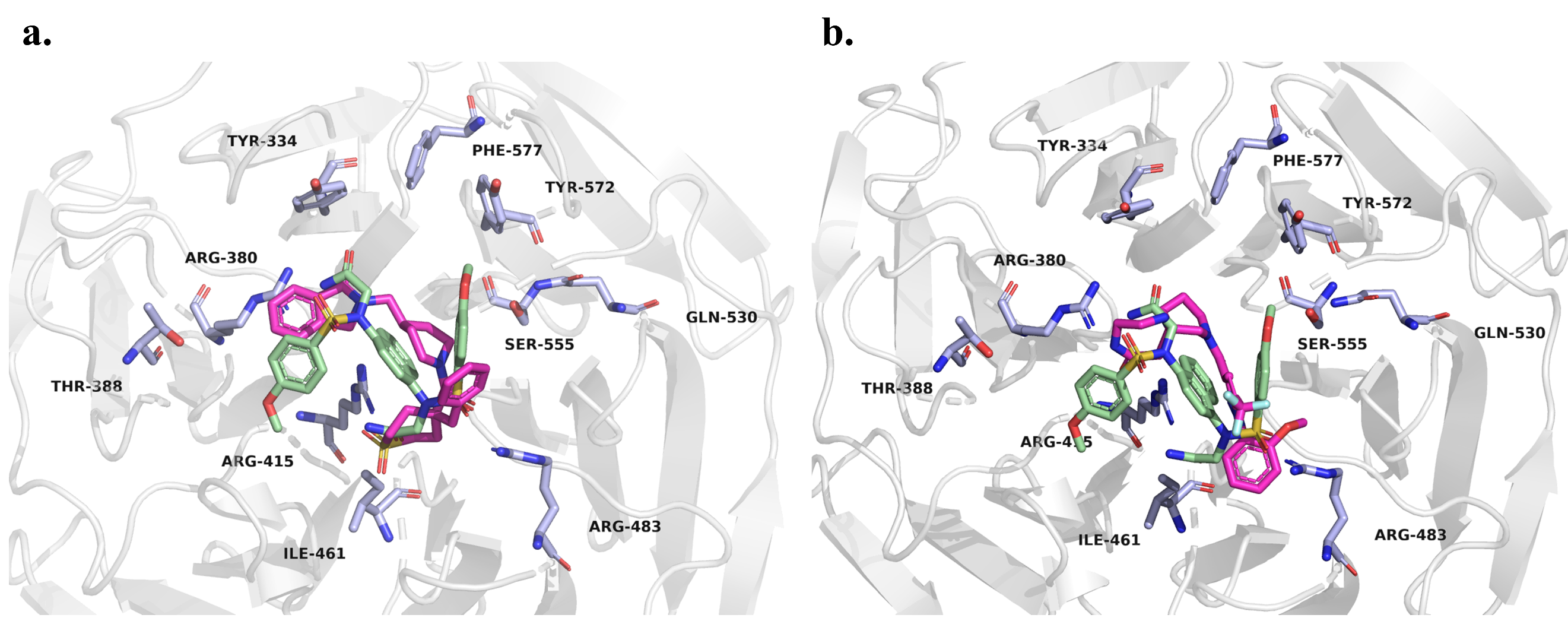

**Figure S2.** (a) Overlap of binding pose of the Keap1 protein with the reference compound (PDB ID: 4XMB) and binding poses of the hit compounds using docking. (pale green: reference compound; magenta: hit1 compound); (b) Overlap of binding pose of the Keap1 protein with the reference compound (PDB ID: 4XMB) and binding poses of the hit compounds using docking. (pale green: reference compound; magenta: hit2 compound).

**Table S1.** List of physiochemical properties of compounds used by MultiPPIMI.

| **Property** | **Description** |
| --- | --- |
| NumRings | Number of rings for a molecule |
| NumAromaticCarbocycles | Number of aromatic carbocycles for a molecule |
| NumAromaticRings | Number of aromatic rings for a molecule |
| NumAliphaticRings | Number of aliphatic (containing at least one non-aromatic bond) rings for a molecule |
| NumAromaticHeterocycles | Number of aromatic heterocycles for a molecule |
| NumHeteroatoms | Number of heteroatoms for a molecule |
| NumSaturatedHeterocycles | Number of saturated heterocycles for a molecule |
| NumSaturatedCarbocycles | Number of saturated carbocycles for a molecule |
| NumSaturatedRings | Number of saturated rings for a molecule |
| NOCount | Number of Nitrogens and Oxygens |

**Table S2.** List of physiochemical properties of proteins used by MultiPPIMI.

| **Property** | **Description** |
| --- | --- |
| PCP_PC | Composition of positively charged residues |
| PCP_NC | Composition of negatively charged residues |
| PCP_NE | Composition of neutral charged residues |
| PCP_PO | Composition of polar residues |
| PCP_NP | Composition of non-polar residues |
| PCP_AL | Composition of residues having aliphatic side chain |
| PCP_CY | Composition of residues having cyclic side chain |
| PCP_AR | Composition of aromatic residues |
| PCP_AC | Composition of acidic residues |
| PCP_BS | Composition of basic residues |
| PCP_NE_pH | Composition of neutral residues based on pH |
| PCP_HB | Composition of hydrophobic residues |
| PCP_HL | Composition of hydrophilic residues |
| PCP_NT | Composition of neutral residues |
| PCP_HX | Composition of hydroxylic residues |
| PCP_SC | Composition of residues having sulphur content |
| PCP_TN | Composition of tiny residues |
| PCP_SM | Composition of small residues |
| PCP_LR | Composition of large residues |

**Table S3**. Hyper-parameters of the baseline models.

| Classifier | Hyper-parameters | Search space | Optimal value |
| --- | --- | --- | --- |
| RF | n_estimators | [50, 100, 150, 200] | 100 |
| SVM | C | [0.1, 0.5, 1.0] | 1.0 |
|  | kernel | [‘linear’, ‘linear’, ‘poly’, ‘rbf’, ‘sigmoid’] | ‘poly’ |
| XGBoost | learning_rate | [0.0001:0.1] | 0.005 |
|  | n_estimators | [50, 100, 150, 200] | 200 |
| MLP | hidden_layer_sizes | [50, 100, 300, 500] | 500 |
|  | learning_rate | [0.0001:0.1] | 0.001 |
|  | batch_size | [125, 256, 512] | 125 |
